# Antimicrobial peptide glatiramer acetate targets *Pseudomonas aeruginosa* lipopolysaccharides to breach membranes without altering lipopolysaccharide modification

**DOI:** 10.1101/2023.05.23.541429

**Authors:** Ronan A. Murphy, Jade Pizzato, Leah Cuthbertson, Akshay Sabnis, Andrew Edwards, Laura M. Nolan, Thomas Vorup-Jensen, Gerald Larrouy-Maumus, Jane C. Davies

## Abstract

Antimicrobial peptides (AMPs) are key components of innate immunity across all kingdoms of life. Both natural and synthetic AMPs are receiving renewed attention in the efforts to combat the antimicrobial resistance (AMR) crisis and the loss of antibiotic efficacy. The gram-negative pathogen *Pseudomonas aeruginosa* is one of the most concerning infectious bacteria in AMR, particularly in people with cystic fibrosis (CF) where respiratory infections are difficult to eradicate and are associated with increased morbidity and mortality. Cationic AMPs exploit the negative charge of lipopolysaccharides (LPS) on *P. aeruginosa* to bind to and disrupt the bacterial membrane(s) and cause lethal damage. *P. aeruginosa* modifies its LPS, via environmental or genetic factors, to neutralise the charge of the cell and evade AMP killing. Free-LPS is also a component of CF sputum, as is anionic extracellular DNA (eDNA), each of which can bind AMPs by electrostatic interaction. Both free LPS and eDNA also feed into pro-inflammatory cycles. Glatiramer acetate (GA) is a random peptide co-polymer of glycine, lysine, alanine, and tyrosine and used as drug in the treatment of multiple sclerosis (MS); we have previously shown GA to be an AMP which synergises with tobramycin against *P. aeruginosa* from CF, functioning via bacterial membrane disruption. Here, we demonstrate direct binding and sequestration/neutralisation of *P. aeruginosa* LPS in keeping with GA’s ability to disrupt the outer membrane. Binding and neutralisation of eDNA was also seen. At CF-relevant concentrations, however, neither strongly inhibited membrane disruption by GA. Furthermore, in both type strains and clinical CF isolates of *P. aeruginosa*, exposure to GA did not result in increased modification of the Lipid A portion of LPS or in increased expression of genetically encoded systems involved in AMP sensing and LPS modification. With this low selective pressure on *P. aeruginosa* for known AMP resistance mechanisms, the potential to neutralise pro-inflammatory CF sputum components, as well as the previously described enhancement of antibiotic function, GA is a promising candidate for drug repurposing.

## INTRODUCTION

The efficacy of antibiotics in treating bacterial infections is decreasing due to antimicrobial resistance (AMR) and the costs continue to mount globally in morbidity and mortality ^1–5^. New tools and strategies are required to deal with increasing AMR, particularly in the absence of development and approval of new antibiotics ^6–9^. *Pseudomonas aeruginosa* is of particular concern in AMR due to its innate, acquired and adaptive resistance mechanisms. It is considered by the World Health Organization (WHO) to be of Critical priority for development of new antimicrobials ^10^. *P. aeruginosa* is a ubiquitous environmental gram-negative bacterium which infects opportunistically across a variety of bodily sites particularly when immunity to infection has been compromised ^11^.

*P. aeruginosa* is especially associated with infections in the lungs of people with cystic fibrosis (CF) where it results in both acute and chronic infections and results in worse long-term outcomes ^12–16^. Antibiotic treatments of infections in CF pose their own, unique set of challenges, on top of those of AMR, including the physical barrier of the characteristic dehydrated CF airway surface liquid (ASL), an acidified environment and the recalcitrant lifestyle of the bacterial biofilm ^17–19^.

In response to the demands of the AMR crisis, attention has increasingly turned to antimicrobial peptides (AMPs); short, usually cationic and amphipathic peptides which occur naturally across the kingdoms of life as a central part of innate immunity ^20,21^. AMPs mostly function as bacterial membrane disruptors, weakening or breaching the bacterial cell envelope (CE) and causing membrane collapse and cell death. The ‘antibiotic-of-last-resort’ colistin (CST) is an AMP which has increasingly come to prominence in response to the loss of efficacy of other antibiotic classes ^22^. Many AMPs (both natural and synthetic) are currently in development or in trials as direct acting agents or as antibiotic adjuvants ^8,23^. While few AMPs have made it to the clinic to-date, predominantly for issues of host cell cytotoxicity at antibacterial concentrations, they have potential advantages over other antimicrobial classes in terms of resistance generation which has generally been shown to occur less frequently with AMPs ^24–26^.

Breaching the membrane of gram-negative bacteria is a major challenge for antibiotic treatments; with two membranes and the added protection of lipopolysaccharide (LPS), accessing the cell cytoplasm is extremely difficult for antimicrobials ^27^. Cationic AMPs attack bacterial cells through electrostatic interactions with the CE; the positively charged AMP attaches to the negatively charged membrane, often via the outermost aspect of the cell, the LPS in the case of *P. aeruginosa* ^21,28,29^. As a defence, *P. aeruginosa* strains can modify their LPS structures by addition of positive charges, neutralising the charge of the cell and reducing AMP binding affinity ^30^. This can take the form of exploiting cations in their environment (preferentially divalent cations Mg^2+^ and Ca^2+^) or by genetically encoded mechanisms ^31,32^. The best studied of these modifications is the addition of 4-amino-4-deoxy-L-arabinose (L-Ara4N) to the Lipid A portion of LPS which is carried out by the Arn operon (ArnBCADTEF) and mediated by a series of two component systems (TCSs) of *P. aeruginosa* (PhoPQ, PmrAB, CprSR, ParSR and ColSR) ^33^. Mutations in these TCSs lead to AMP resistance with true CST resistance resulting from changes which deactivate TCS sensor gene(s) leading to constitutive Arn operon expression and LPS modification ^32,34–38^. As well as L-Ara4N, other encoded Lipid A modification types include additions of phosphate, C10:3OH species, palmitate and combinations of these with L-Ara4N- and palmitate-modified LPS being associated with increased airway disease severity ^32,39^.

Free-LPS (shed by bacteria and from dead bacteria) is a significant component of the sputum of people with CF, as is extracellular DNA (eDNA), with both having been shown to have detrimental effects on the host ^40,41^. LPS is highly pro-inflammatory and its presence in the CF lung contributes to cycles of inflammation seen in that environment and eDNA is a vital component of bacterial biofilms, promoting biofilm development and bacterial recalcitrance. Both LPS and eDNA feed into cycles of infection and inflammation in CF ^42–45^. Presence of both may also have consequences for AMP activity in the CF lung, as they are negatively charged and capable of binding and sequestering cationic AMPs as well as competing for divalent cations ^46,47^. Conversely, the affinity of AMPs for LPS has been proposed as being beneficial, leading to sequestration and neutralisation of free-LPS thereby limiting its pro-inflammatory capacity ^48,49^.

We have previously shown *in vitro* that the multiple sclerosis (MS) drug glatiramer acetate (GA) also functions as an AMP and synergises with the aminoglycoside antibiotic, tobramycin, against *P. aeruginosa* from people with CF ^50,51^. That work also showed GA is a bacterial membrane disruptor, in common with many AMPs ^21^. Produced by the random polymerisation of the four N-carboxy-α-amino acid anhydrides of L-glutamate, L-lysine, L-alanine, and L-tyrosine, in MS GA functions as an immunomodulator with much of its known activity shown to be anti-inflammatory in that condition ^52–55^.

Having previously demonstrated GA’s ability to permeabilise *P. aeruginosa* cells and breach the CE while sensitising them to antibiotic treatment, here, we examine whether GA interacts with *P. aeruginosa* LPS via electrostatic interaction as its point of contact with the bacterial cell in common with other cationic AMPs. With the known interplay at a variety of levels between AMPs, LPS, divalent cations and eDNA and the presence and importance of each in the lungs of people with CF, it is necessary to understand their implications for GA activity in the CF airway. We also tested whether exposure to GA resulted in *P. aeruginosa* mounting a defensive response in the manner seen for other AMPs with modification of LPS and Lipid A. With a proposed benefit of AMPs being the decreased likelihood with which they generate resistance, it is important to know if GA is stimulatory to the TCSs commonly associated with AMP protection and LPS modification and potentially selective for resistance.

## MATERIALS AND METHODS

### Strains and growth conditions

*P. aeruginosa* type strains PAO1, PA14 and PAK and 11 clinical *P. aeruginosa* isolates were used in this study. The panel of clinical strains, which was assembled for a previous study, constitutes isolates from the CF Bacterial Repository at the National Heart and Lung Institute, Imperial College London from airway samples of people with CF at the Royal Brompton Hospital, London (**Supplementary Table 1**) ^51^. Bacteria were stored in Microbank vials (Pro-Lab Diagnostics) at -80°C. Isolates were grown overnight at 37°C on LB agar (Merck). Single colonies were inoculated into Mueller-Hinton broth (MHB) (Merck) and incubated overnight at 37°C with agitation at 200 r.p.m.

### LPS quantification

LPS was quantified as Endotoxin Units (EU) using a limulus amoebocyte lyase (LAL) Endotoxin Quant Kit (ThermoFisher), per manufacturer’s instructions. *P. aeruginosa* LPS (Merck) at physiologically relevant CF concentrations of 0.01 and 0.02 mg/mL and the supraphysiological concentration of 0.1 mg/mL was incubated at 37°C for 30 mins, with and without 50 mg/L GA (Biocon), as was GA without LPS ^39,56,57^. Samples were diluted 1:1000 in in phosphate buffered saline (PBS), to bring them within the sensitivity range of the kit, before addition to the remaining kit components along with blanks and standards. Each condition was tested in triplicate. The optical density was measured at 405 nm (OD_405_) in FLUOstar Omega platereader (BMG Labtech). Blanked Standards were fitted using simple linear regression to create a standard curve (GraphPad Prism). Concentrations of samples in EU were interpolated from the standard curve via their blanked OD_405_ results. Neutralisation of LPS by GA was calculated as the EU of LPS in the presence of GA as a percentage of the EU of LPS in the absence of GA, at each LPS concentration tested, and compared to No Neutralisation (i.e. 0%).

### DNA extraction and eDNA quantification

Bacterial DNA was extracted from strains PAO1, PA14 and PAK from ∼1 × 10^8^ CFU using NucleoSpin Microbial DNA Mini kit (Macherey-Nagel), as per manufacturer’s instructions. Purified DNA was stored at -20°C. DNA concentrations were measured on a NanoDrop (ThermoFisher) and adjusted to 100 mg/mL. DNA from the individual strains were pooled together for use as eDNA and diluted to required final concentrations.

Concentrations of eDNA were chosen as a CF physiologically relevant concentration (1 mg/mL) and a supraphysiological concentration (10 mg/mL) ^58^. Each eDNA concentration was combined with GA at concentrations of 0, 6.25, 12.5, 25, 50 and 100 mg/L and incubated at 37°C for 30 mins, as were eDNA-free GA solutions. Propidium iodide (PI) was added to the suspensions to a concentration of 1 µg/mL and fluorescence measured in a platereader at excitation 544 nm, emission 610 nm, in the wells of a black microtitre plate (ThermoFisher) (5 wells of each). Neutralisation of eDNA by GA was calculated as the fluorescence intensity of eDNA in the presence of GA as a percentage of the fluorescence intensity of eDNA in the absence of GA, at each eDNA concentration tested. Each was compared to No Neutralisation at the same eDNA concentration (i.e. 0 %).

### Conditions for membrane disruption assays

The effects of *P. aeruginosa* LPS on the previously reported rapid activity of GA against *P. aeruginosa* membranes were tested in two different modes. Firstly, a ‘Pre-incubation’ was performed to test the ability of LPS to sequester GA where LPS and GA were incubated together at 37°C for 30 mins before administration to cell suspensions. Secondly, in the ‘Background’ of the assays, to test the effect of LPS presence on administered GA activity, LPS was added to each assay buffer. LPS-only and GA-only solutions were also incubated under the same conditions. In both cases final concentrations of *P. aeruginosa* LPS of 0.01, 0.02 and 0.1 mg/mL and 50 mg/L GA were used (as above). To further test for GA-LPS interactions, the protective effect of Mg^2+^ cations were added to assay buffers as MgSO_4_ (Merck) at CF physiologically relevant levels (0.5 and 1 mM) ^59,60^. Finally, to test for GA sequestration by eDNA, 50 mg/L GA and 1 mg/mL eDNA were incubated together at 37°C for 30 mins before administration to cell suspensions. eDNA-only and GA-only solutions were also incubated under the same conditions. Assay specific buffers (see below) were used in each case.

### Outer membrane disruption

Disruption of the bacterial outer membrane (OM) of *P. aeruginosa* was measured using the fluorescent probe 1-*N*-Phenylnaphthylamine (NPN) (Merck) at a final concentration 10 µM ^61^. Late-exponential phase cultures of *P. aeruginosa* were washed twice with 5 mM HEPES and adjusted to OD_600_ 0.5 in appropriate buffer depending on whether a Background or Pre-incubation experiment was required. Conditions were tested as outlined above. Controls included dye free wells, GA-only treated bacteria, wells without GA treatments but containing LPS/eDNA/Mg^2+^. Triplicate technical repeats were carried out and averaged for each biological replicate performed. Assays were performed in black microtitre plates (ThermoFisher) in 200 µL volumes and fluorescence was measured in a platereader at excitation 355 nm, emission 460 nm every 30 secs for 15 mins. Mean NPN Uptake Factor over 15 mins was calculated as ([Fluorescence of Sample with NPN – Fluorescence of Sample without NPN] / [Fluorescence of buffer with NPN – Fluorescence of buffer without NPN]). At least three biological replicates were performed on each *P. aeruginosa* type strain. Comparisons to GA activity were performed separately for each element tested (LPS or divalent cations or eDNA).

### Cytoplasmic membrane depolarisation

Depolarisation of the cytoplasmic membrane (CM) of *P. aeruginosa* isolates was measured using the fluorescent probe 3,3’-Dipropylthiadicarbocyanine Iodide (DiSC_3_(5)) (Thermo Scientific) ^62^. Late-exponential phase cultures of *P. aeruginosa* were washed twice and adjusted to an OD_600_ of 0.05 in 5 mM HEPES-20 mM glucose. DiSC_3_(5) was added to the bacterial cultures to a concentration of 1 µM and aliquoted to the wells of a black microtitre plate in 200 µL volumes with triplicate technical repeats. The fluorescent signal of the dye was allowed to quench for 30 mins in the dark before addition of test solutions. Conditions were tested as outlined above. Controls included dye free wells, GA-only treated bacteria, wells without GA treatments but containing LPS/eDNA/Mg^2+^. Fluorescence was measured in a platereader at excitation 544 nm, emission 620 nm every 30 secs for 15 mins. Fluorescent signals of samples were normalised to a cell free background with identical components and averaged. At least three biological replicates were performed on each *P. aeruginosa* type strain. Comparisons to GA activity were performed separately for each element tested (LPS or divalent cations or eDNA).

### Cell envelope permeability

Permeabilisation of the *P. aeruginosa* CE was measured using the fluorescent dye PI as per manufacturer’s instructions (Merck, UK). Late-exponential phase cultures of *P. aeruginosa* were washed twice and adjusted to an OD_600_ of 0.5 in PBS. PI was added to the bacterial cultures to a concentration of 1 µg/mL. Conditions were tested as outlined above. Fluorescence was measured in a platereader at excitation 544 nm, emission 610 nm every 30 secs for 1 hr in the wells of a black microtitre plate in 200 µL volumes. Technical triplicates were performed, blanked and signal averaged in each biological experiment. Area under the curve (AUC) of the PI fluorescence intensity was computed using the trapezoid rule over 1 hr (GraphPad Prism). At least three biological replicates were performed on each *P. aeruginosa* type strain. Comparisons to GA activity were performed separately for each element tested (LPS or divalent cations or eDNA).

### AMP exposure

Overnight cultures of *P. aeruginosa* grown in MHB were centrifuged (3500 *g*, 15 mins), supernatant discarded, bacterial pellets resuspended in fresh media and cultures adjusted to OD_600_ 0.05 (5 × 10^6^ CFU/mL) in MHB. AMPs were added to *P. aeruginosa* type strain cultures at the following final concentrations; 50 mg/L GA, 0.5 mg/L CST or 16 mg/L LL-37, along with untreated cultures. Clinical strains were incubated with No Treatment or 50 mg/L GA. After AMP addition, cultures were incubated at 37°C with agitation at 200 r.p.m. and bacteria allowed to grow to mid-log phase (∼4 hrs). Triplicate biological replicates of each strain at each condition were performed with the exception of LL-37 exposure, which was performed in duplicate due to resource limitation.

After incubation, OD_600_ was measured and ∼1 × 10^8^ CFU were harvested and placed in a 1.5 mL eppendorf and centrifuged at 15000 *g* for 10 mins. The supernatant was removed and the pellet was used for RNA extraction and gene expression (details below). The remainder of each culture was centrifuged (3500 *g*, 15 mins), supernatant discarded and pellet resuspended in 1 mL sterile PBS used for MALDI-TOF mass spec analysis of Lipid A (details below).

### Gene expression

Bacterial RNA was extracted from ∼1 × 10^8^ CFU after AMP exposure (above) using Direct-zol RNA Miniprep kit (Zymo) as per manufacturer’s instructions. Expression levels of TCS response regulator genes *phoP*, *pmrA*, *cprR*, *parR*, Arn operon gene *arnB* and housekeeping gene *rpsL* were measured using primers from Lee *et al.* (2014) (ThermoFisher) ^34^. Quantitative Real-Time PCR was performed using KAPA SYBR FAST One-Step kit (KAPA) on a QuantStudio 7 Flex (ThermoFisher) with the following cycling conditions; 5 mins at 42°C for Reverse Transcription, 3 mins at 95°C for Enzyme Activation followed by 40 cycles of 10 secs at 95°C (Denaturation) and 30 secs at 60°C (Annealing/Extension). Triplicate wells of each PCR were performed as technical replicates. Cycle thresholds (Ct) for each well were calculated by the QuantStudio analysis software (ThermoFisher) and averaged. Expression of each gene was normailised to the expression of *rpsL* (ΔCt) and relative expression of each gene under AMP stimulation was normailised to its expression under No Treatment (ΔΔCt).

### MALDI-TOF

Analysis of Lipid A conformations were carried out as previously described ^63^. Briefly, bacteria were centrifuged at 17000 × *g* for 2 mins, supernatant discarded and the pellet was washed three times with 300 μL of ultrapure water and resuspended to a density of McFarland 20 as measured using a McFarland Tube Densitometer followed by acetic acid hydrolysis 1% at final concentration at 98°C for 1 hour. After 2 washes with ultrapure water, the hydrolysate was suspended in 50 μL and a volume of 0.4 μL of this suspension was loaded onto the MALDI target plate overlaid with 1.2 μL of Norharmane matrix (Sigma-Aldrich) solubilised in chloroform/methanol (90:10 v/v) to a final concentration of 10 mg/mL.

The samples were loaded onto a disposable MSP 96 target polished steel BC (Bruker Part-No. 8280800). The bacterial suspension and matrix were mixed directly on the target by pipetting. The spectra were recorded in the linear negative-ion mode (laser intensity 95 %, ion source 1 = 10.00 kV, ion source 2 = 8.98 kV, lens = 3.00 kV, detector voltage = 2652 V, pulsed ion extraction = 150 ns). Each spectrum corresponded to ion accumulation of 5,000 laser shots randomly distributed on the spot. The spectra obtained were processed with default parameters using FlexAnalysis v.3.4 software (Bruker Daltonik).

For each biological replicate performed, the abundance of native and modified Lipid A were enumerated via the AUCs of their spectra at each mass to charge ratio (*m*/*z*) where each appears (**Supplementary Table 2**). Using the total values across all *m*/*z* where each modification is seen, the ratio Native:Modified was calculated for each modification type (phosphate, C10:3OH, palmitate and L-Ara4N). One biological replicate of strain GA899 was lost during processing and subsequent data for this strain is therefore for biological duplicates.

### Genomics

DNA extractions for genomic sequencing were performed using Maxwell™ RSC Cell DNA Purification Kit and automated extraction on the Maxwell RSC 48 Instrument (Promega). The DNA concentration was evaluated using a Qubit dsDNA BR assay kit and Qubit Fluorometer (ThermoFisher). Sequencing was on a standard Illumina HiSeq platform.

Genomes were assembled with Shovill and annotated using Prokka with quality control by Quast on the Galaxy platform ^64–72^. The sequences of the genes of two component systems PhoPQ, PmrAB, CprSR, ParSR and ColSR of PAO1 were taken from https://pseudomonas.com/ and used to extract their corresponding sequences from the genomic data of the 11 clinical *P. aeruginosa* using the MyDbFinder tool (https://cge.food.dtu.dk/services/MyDbFinder/) ^73^. Gene sequences from clinical strains were aligned with those from PAO1 using MEGA X and non-synonymous SNPs identified ^74^. Gene accession numbers can be found in Supplementary Table 3.

### Statistics

Unpaired, non-parametric data was analysed using analysis of variation (ANOVA) Kruskal-Wallis method with Dunn’s multiple correction for non-parametric (adjusted p values reported). Membrane assay data biological replicates were log transformed and compared with Welch’s ANOVA with Dunnett’s T3 for multiple comparison and with Welch’s t test for two datasets. Paired comparisons of multiple non-parametric datasets used Friedman ANOVA test method, with Dunn’s multiple correction (adjusted p values reported). Paired comparisons of two non-parametric datasets used Wilcoxon tests. A significant difference was reported with p < 0.05. Biological replicates comprise means of technical replicates. All analyses and data presentation were performed in GraphPad Prism version 9.0 or later.

## RESULTS

### Bilateral sequestration of GA and *P. aeruginosa* LPS

We first tested whether GA interacted directly with *P. aeruginosa* LPS as a potential mechanism of action of the drug. A known active concentration of GA (50 mg/L) was incubated with concentrations of LPS at supraphysiological level (0.1 mg/mL) and two concentrations relevant to the CF lung environment (0.02 and 0.01 mg/mL) at 37°C for 30 mins ^39,50,51,56,57^. We determined the availability of free LPS with an Endotoxin detection kit reasoning that if GA targets LPS via electrostatic interactions the resulting GA-LPS complex would reduce the availability of LPS for quantification, as has been shown for other AMPs ^48,75–77^. GA significantly neutralised LPS at concentrations of 0.02 (56.9 ± 4.7 %) and 0.01 mg/mL (50.7 ± 12.8 %) (p < 0.05) indicating direct binding of GA to LPS (**Figure 1A**).

**Figure 1.**
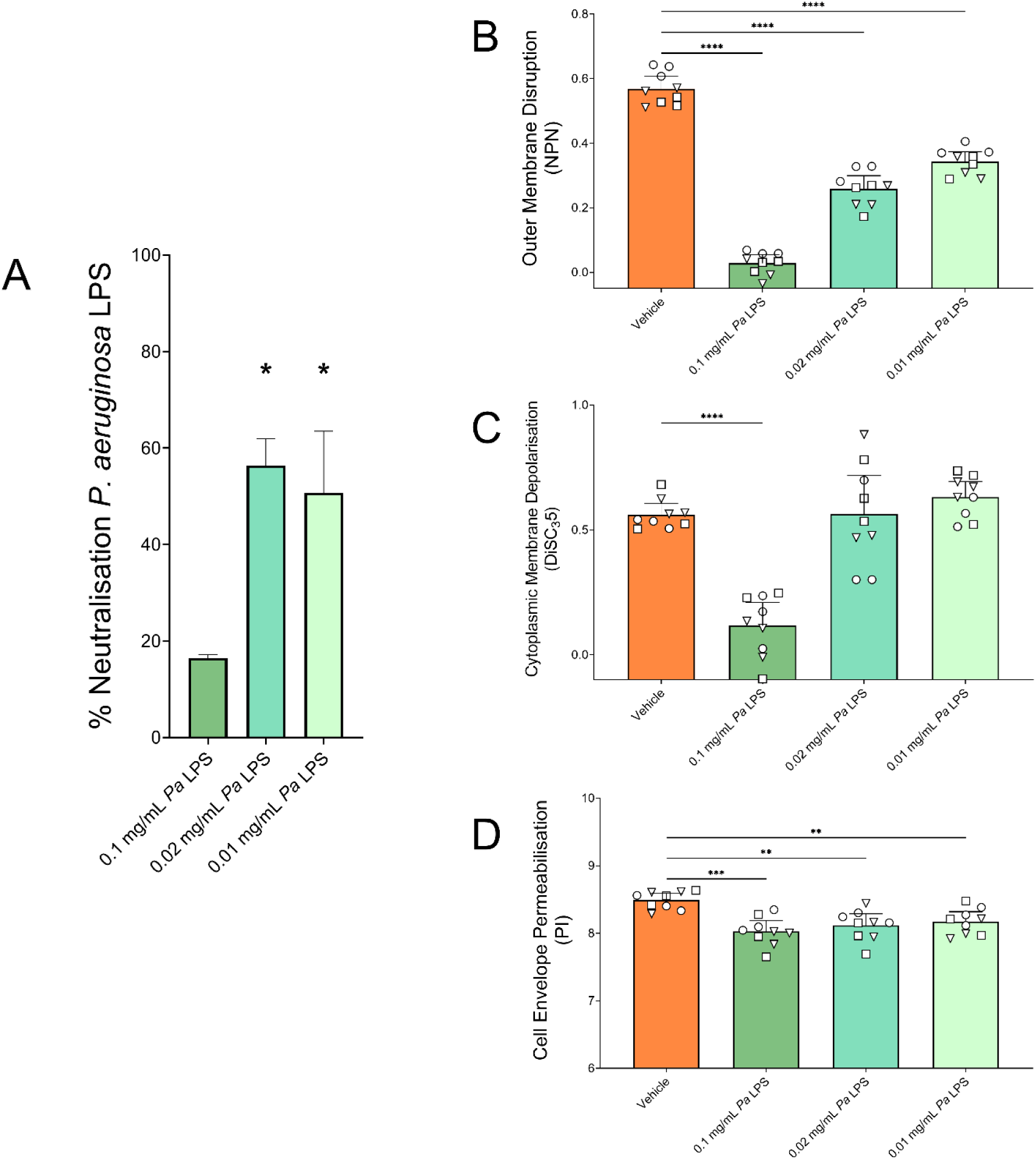
Bilateral sequestration of GA and P. aeruginosa LPS. **A**. Neutralisation of P. aeruginosa LPS was calculated as the percentage reduction of the quantifiable LPS in the presence of GA at each LPS concentration. Incubation of 0.02 (p = 0.0186) and 0.01 mg/mL (p = 0.0499) LPS with GA significantly increased neutralisation of LPS, compared to no GA (Kruskal-Wallis test with Dunn’s multiple comparison). (Median with 95%CI. n = 3). **B**. The ability of 50 mg/L GA to disrupt the Outer Membrane of P. aeruginosa type strains was significantly reduced by 30 mins pre-incubation with all P. aeruginosa LPS concentrations tested (each p < 0.0001). **C**. The ability of 50 mg/L GA to depolarise the Cytoplasmic Membrane of P. aeruginosa type strains was significantly reduced by incubation with P. aeruginosa LPS concentration of 0.1 mg/mL (p < 0.0001) but not 0.02 mg/mL or 0.01mg/mL. **D**. The ability of 50 mg/L GA to permeabilise the Cell Envelope of P. aeruginosa type strains was significantly reduced by incubation with all P. aeruginosa LPS concentrations tested – 0.1 (p = 0.0002), 0.02 (p = 0.0024) and 0.01 mg/mL (p = 0.0031). B-D Log transformed data tested using Welch ANOVA with Dunnett’s T3 multiple comparison. Medians with 95% CIs of biological replicates (n = 9) of P. aeruginosa PAO1 (ϒ), PA14 (♦) or PAK (ρ).

The converse, sequestration of GA by LPS, was tested for using GA’s known membrane disruption properties as a proxy, due to the difficulties in accurately assaying for a highly heterogeneous peptide such as GA which can take on > 10^30^ different peptide chains ^51^. We assessed the impact on the outer membrane (OM), cytoplasmic membrane (CM) and on the overall cell envelope (CE) permeability using *P. aeruginosa* LPS at the same concentrations: supraphysiological (0.1 mg/mL) and CF-lung-relevant (0.02 and 0.01 mg/mL). Disruption of the OM and permeabilisation of the CE of *P. aeruginosa* type strains by GA was significantly reduced after GA had been pre-incubated (37°C for 30 mins) with LPS (p < 0.01) (**Figure 1A, C**). The highest concentration of LPS tested also significantly reduced the ability of GA to depolarise the bacterial CMs (p < 0.0001) (**Figure 1B**). These results indicate that available concentrations of both LPS and GA are reduced by presence of other and that sequestration is bilateral.

### Membrane perturbing activity of glatiramer acetate in the presence of CF-physiological *P. aeruginosa* LPS and divalent cations concentrations

With the evidence of direct interactions between GA and LPS, and the impact that pre-incubation had on GA’s action on bacterial cell membranes, we next tested the ability of GA to function if *P. aeruginosa* LPS was present in the background, as it will be in the CF lung environment. To do this, the previously reported membrane perturbation properties of GA were examined with LPS in the assay buffer, to test the effect of LPS presence on the known rapid action of GA on bacterial membranes. *P. aeruginosa* LPS significantly reduced the OM disruption of strains PAO1, PA14 and PAK at each concentration tested and the highest concentration also reduced CM depolarisation (p ≤ 0.05) (**Figure 2A, B**). No effect was seen on the ability of GA to permeabilise the CEs at the LPS concentrations tested (**Figure 2C**).

**Figure 2.**
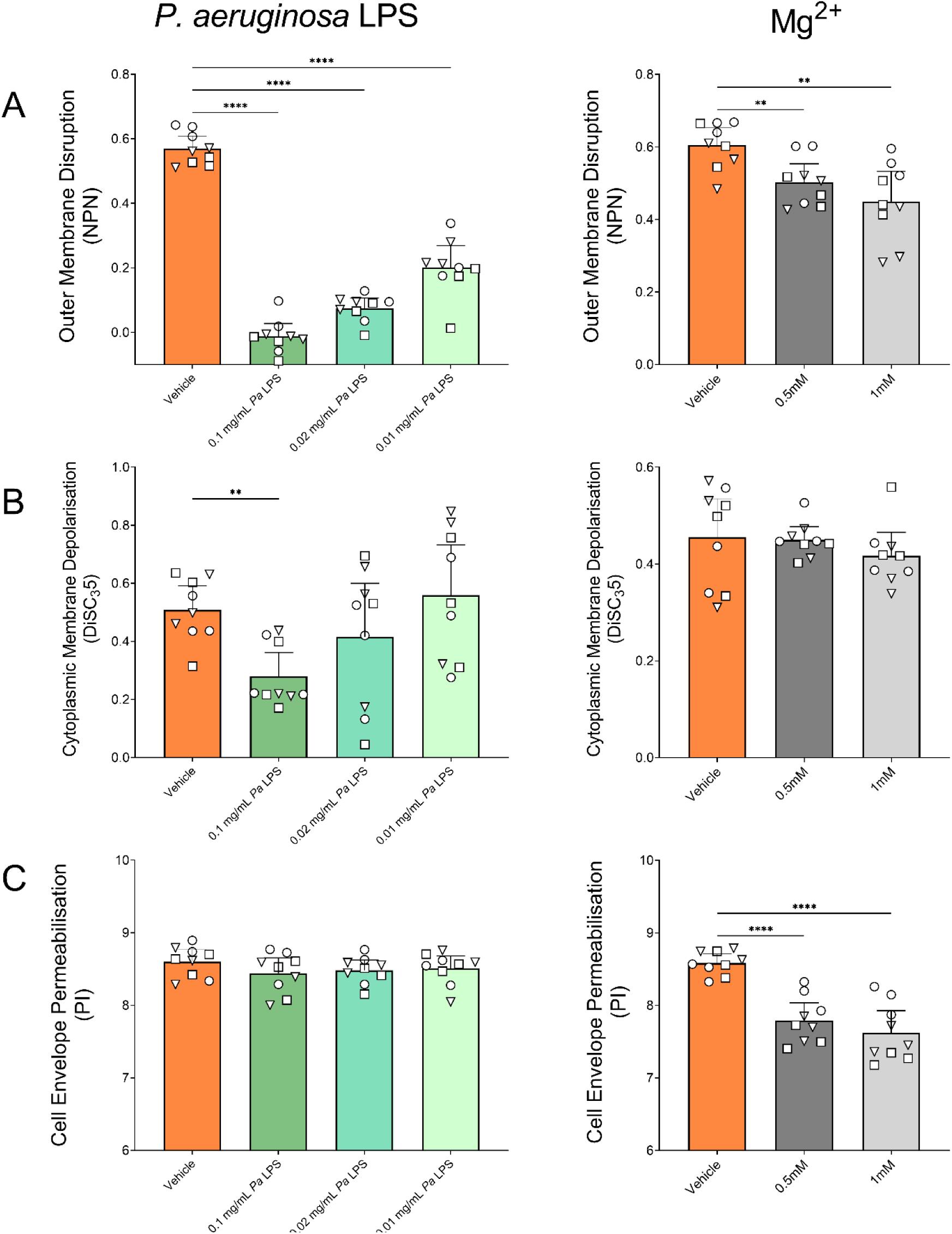
Effect of presence of CF sputum components on membrane perturbations resulting from 50 mg/L GA in P. aeruginosa strains PAO1, PA14 and PAK. **A**. Disruption of the OM of P. aeruginosa by GA was significantly reduced by each concentration of LPS tested (each p < 0.0001) while Mg^2+^ at 1 (p = 0.0051) and 0.5 mM (p = 0.0081) both also reduced GA activity. **B**. Depolarisation of the CMs of P. aeruginosa by GA was only significantly reduced by LPS at the supraphysiological concentration of 0.1 mg/mL (p = 0.001) and was unaltered by Mg^2+^ presence. **C**. Permeabilisation of the CE of P. aeruginosa by GA was not seen for any LPS concentration tested but was significantly reduced by Mg^2+^(each p < 0.0001). Log transformed data tested using Welch ANOVA with Dunnett’s T3 multiple comparison. Medians with 95% CIs of biological replicates (n = 9) of P. aeruginosa PAO1 (ϒ), PA14 (♦) or PAK (ρ).

The protective properties of Mg^2+^ against GA activity were also tested at physiological concentrations at mid - (0.5 mM) and upper-levels (1 mM) ^59,60^. Both concentrations of Mg^2+^ were protective against GA OM disruption and CE permeabilisation, with significant decreases in GA activity seen (p < 0.05) (**Figure 2A, C**), but this was not the case for depolarisation of the CM where no change was seen (**Figure 2B**). These results further support the evidence for cellular LPS as a target for GA binding by indicating competitive displacement between the two cations.

### GA binds directly to eDNA

We next wanted to test interactions of GA and eDNA, due to their opposing charges, previous indications of GA-DNA aggregation and the high concentrations of eDNA in CF sputum playing a role in chronic biofilm formation ^50,78^. With both supraphysiological (10 mg/mL) and physiological (1 mg/mL) concentrations of eDNA, GA reduced detectable eDNA using PI fluorescence in a dose-responsive fashion. Thus, as GA concentrations increased, detectable eDNA decreased, reaching statistical significance by 25 mg/L (p < 0.05) (**Figure 3A**). This indicates binding and sequestration of eDNA by GA.

**Figure 3.**
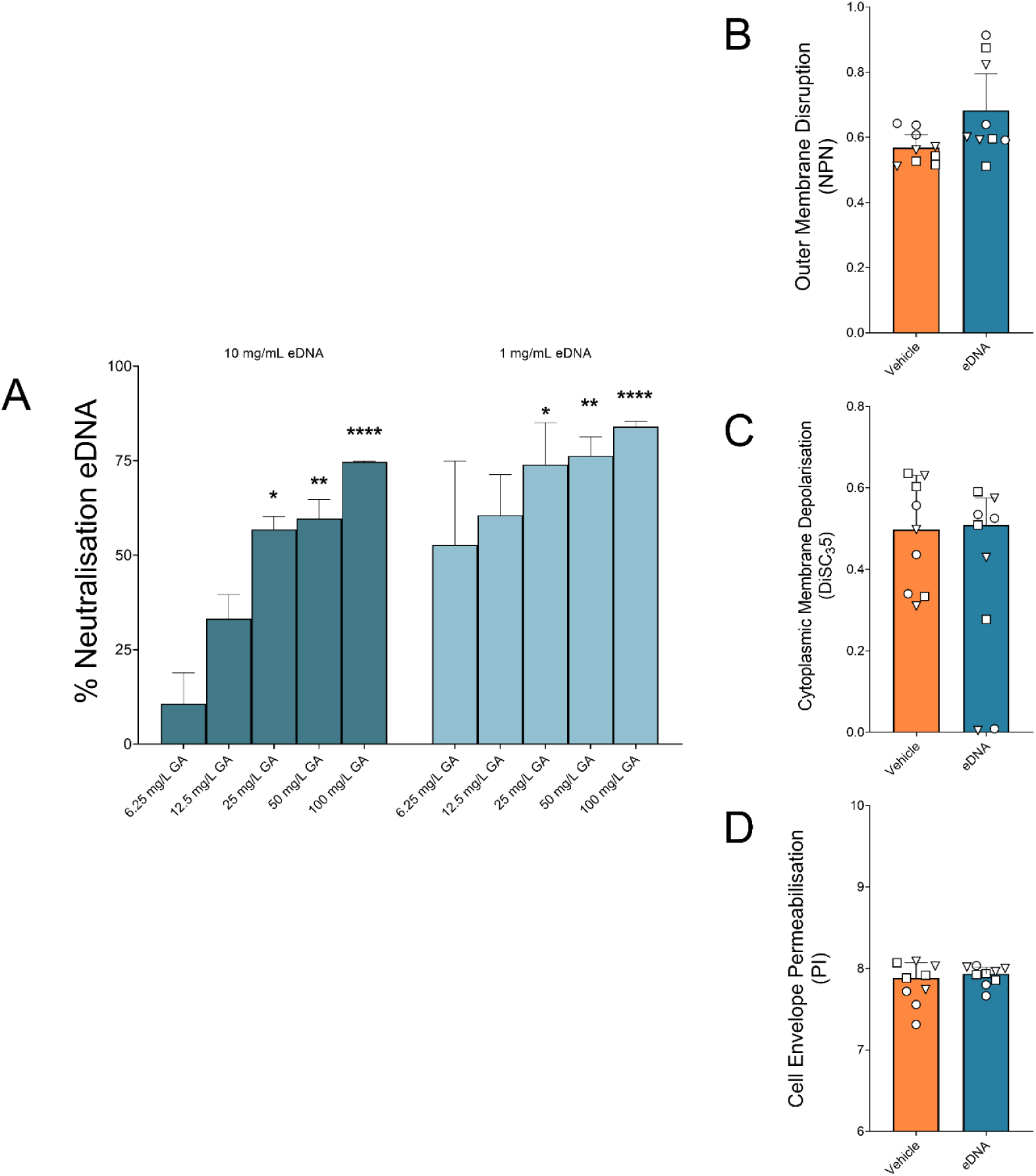
Effect of GA on eDNA availability and effect of CF-relevant eDNA on GA activity. **A**. Neutralisation of eDNA was calculated as the percentage reduction of the detectable eDNA in the presence of GA at each eDNA concentration. All GA concentrations > 25 mg/L significantly increased eDNA neutralisation. At 1 mg/mL eDNA, GA at 25 (p = 0.0111), 50 (p = 0.0068) and 100 mg/mL (p < 0.0001) while at 10 mg/mL eDNA, GA at 25 (p = 0.0223), 50 (p = 0.0027) and 100 mg/mL (p < 0.0001) showed neutralisation (Kruskal-Wallis test with Dunn’s multiple comparison). (Medians with 95%CI. n = 5). The CF-relevant concentration of 1 mg/mL eDNA did not alter the ability of 50 mg/L GA to disrupt the OM (**B**), depolarise the CM (**C**) or permeabilise the CE (**D**) of P. aeruginosa PAO1 (ϒ), PA14 (♦) or PAK (ρ). Welch’s t test of Log transformed data. Medians with 95%CIsof biological replicates (n = 9).

Similarly, to the experiments with LPS, we questioned whether such sequestration occurred bilaterally, next assessing whether eDNA binding, reduced the activity of GA. As with LPS, membrane perturbation assays were employed, after incubating GA-eDNA together at a CF-physiological concentration of eDNA (1 mg/mL). Unlike LPS, no significant differences were seen in the ability of GA to perturb the OM, CM and CE of *P. aeruginosa* strains PAO1, PA14 and PAK in these assays (**Figure 3B-D**).

### Modification of *P. aeruginosa* Lipid A of type strains after GA and AMP exposure

Having identified the interaction of GA and LPS and in the knowledge that many AMPs target the Lipid A component of LPS for membrane attachment, we next tested the reaction of *P. aeruginosa* to the targeting of its LPS by GA. *P. aeruginosa* has been shown to react to attack by other AMPs by sensing the peptides via Two Component Systems (TCSs) and modifying the Lipid A portion of its LPS structures, reducing the charge of its cell envelope and vulnerability to cationic AMPs. MALDI-TOF mass spectrometry was used to investigate if exposure to GA induces changes in the Lipid A structures, as Lipid A modification can result in resistance to AMPs ^79^. The MALDI-TOF spectra of the strains PAO1, PA14 and PAK were acquired with and without exposure to AMPs (GA, CST and LL-37) and abundances of each Lipid A modification type (additions of phosphate, C10:3OH groups, palmitate and L-Ara4N) were normalised to native Lipid A, by ratio. No significant differences were seen in any modification of Lipid A across the *P. aeruginosa* type strains after GA exposure, when compared to the cultures without GA stress (**Figure 4**). LL-37, as a positive inducer of LPS modification, showed the highest (but non-significant) addition of C10-3OH and L-Ara4N in the 3 strains ^80^. No significant changes in LPS Lipid A modification resulted from CST exposure in the CST-sensitive type strains.

**Figure 4.**
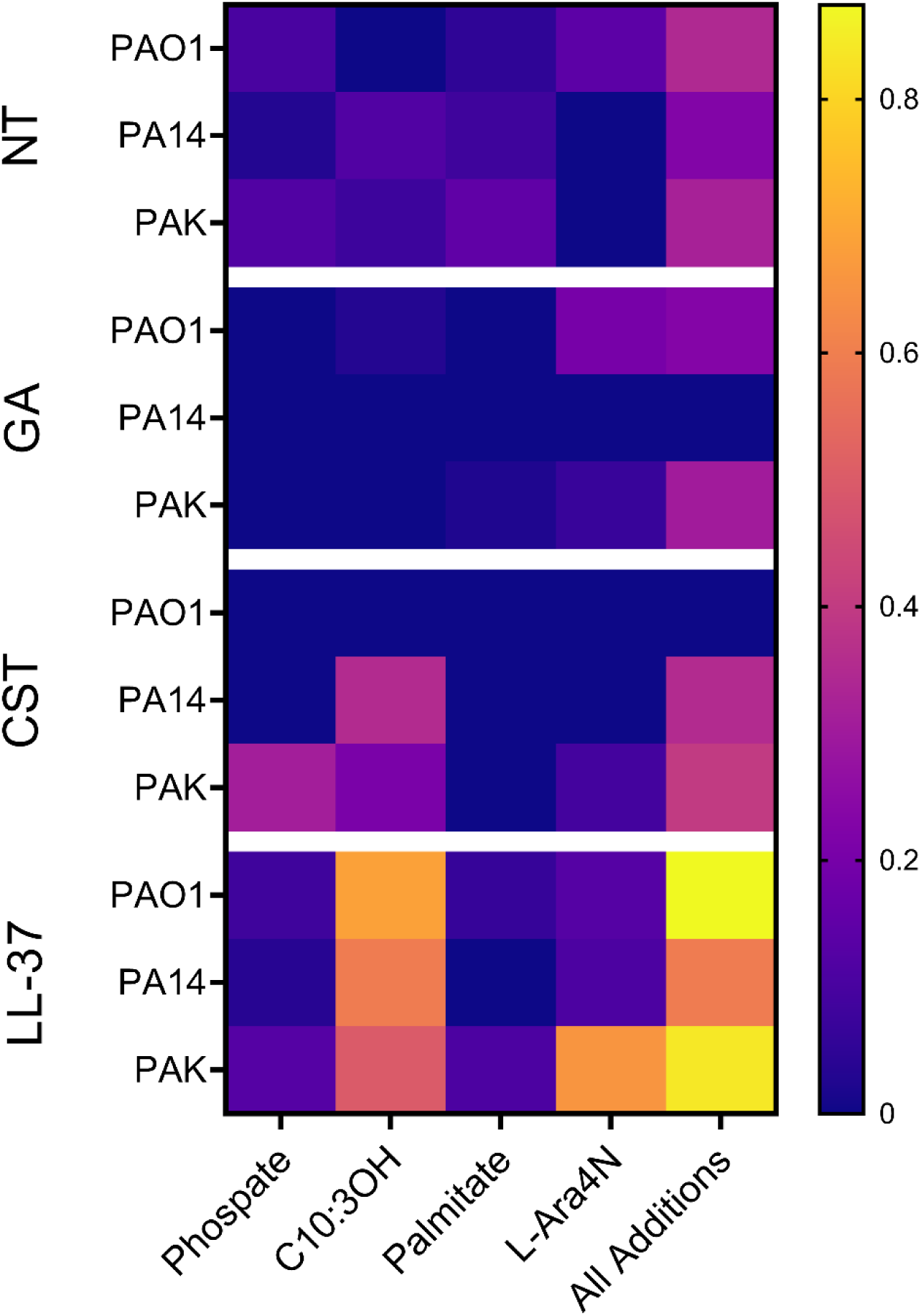
Additions to LPS Lipid A of P. aeruginosa type strains PAO1, PA14 and PAK in response to AMP exposure. Heatmap of ratios of Native Lipid A:Modified Lipid A for each modification type. No significant differences from No Treatment (NT) were seen for any modification type as the result of any AMP tested; 50 mg/L GA, 0.5 mg/L CST or 16 mg/L LL-37 (Friedman test with Dunn’s multiple comparison). Median values of triplicate biological replicates, except LL-37 which was run in duplicate due to resource limitation.

### Modification of *P. aeruginosa* Lipid A of clinical strains from people with CF after GA exposure

We next investigated changes in the Lipid A structures of 11 clinical *P. aeruginosa* isolates from people with CF, after the bacteria were exposed to GA, determining their lipid A profiles with and without treatment with GA and using clinical isolates which had already be characterised for GA-antibiotic synergy ^51^. As previously, each modification type was examined as a ratio with native Lipid A and GA-exposed cultures compared to untreated. Clinical respiratory strains from people with CF showed no significant changes for any modification type of Lipid A (**Figure 5**).

**Figure 5.**
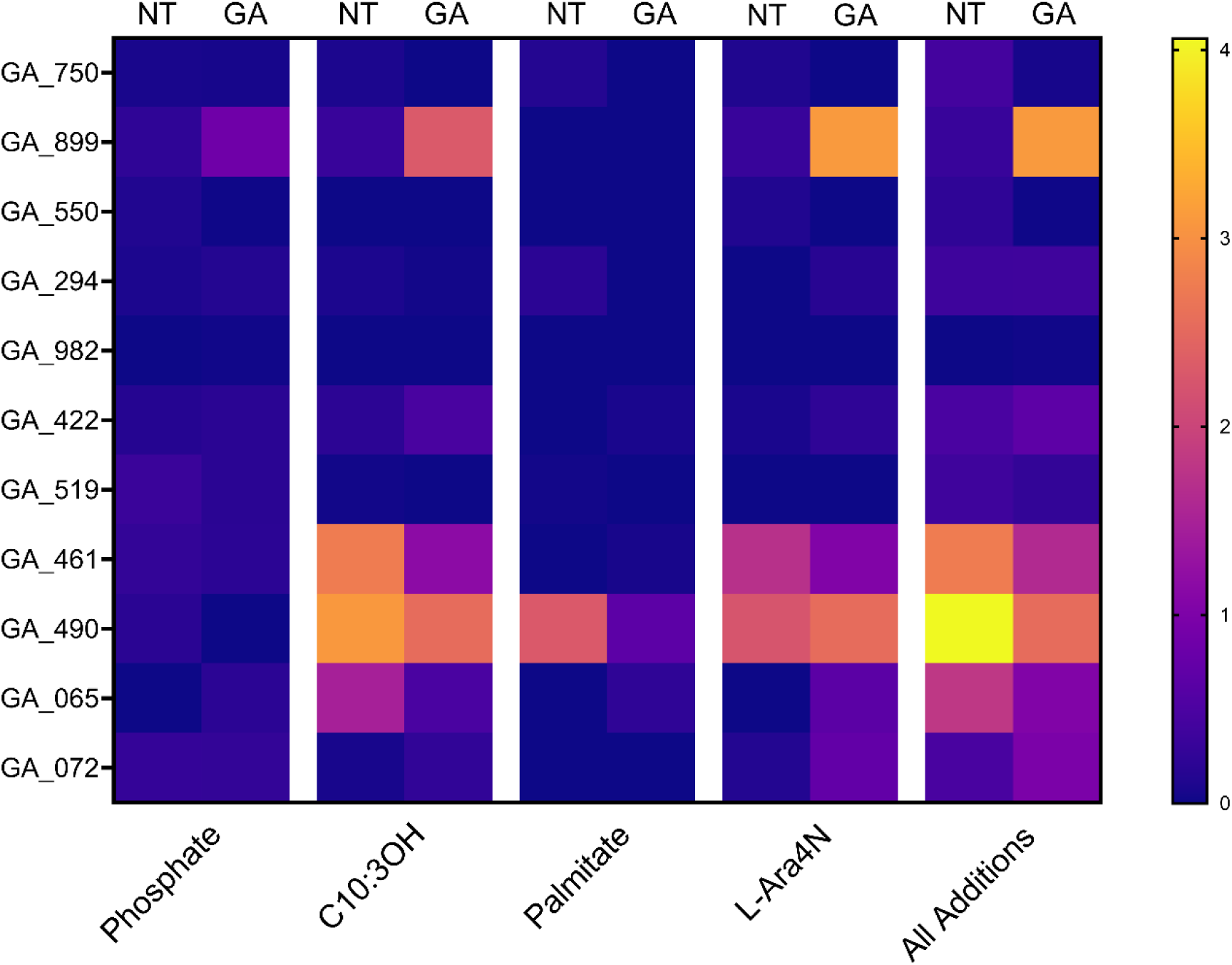
Changes in Lipid A of P. aeruginosa clinical strains from people with CF in response to GA exposure. Heatmap of the ratios of Modified Lipid A:Native Lipid A for each modification type. No significant differences resulted from exposure to 50 mg/L GA, from No Treatment, for any modification type across the 11 clinical strains (Wilcoxon test). Median values of triplicate biological replicates. *GA_899 biological duplicates displayed as one replicate lost for technical reasons. Median value of biological replicates displayed.

### Expression of Two Component Systems (TCS) and *arn* operon genes of *P. aeruginosa* type strains after GA and AMP exposure

In the absence of significant LPS Lipid A modification increases in response to GA exposure and for more detail on the bacterial response to GA, the expression of the response regulator genes of TCSs (*phoP*, *pmrA*, *cprR* and *parR*) frequently associated with AMP detection as well as a gene involved in the LPS L-Ara4N modification process (*arnB*) were investigated using qRT-PCT ^34,81–83^. The effect of exposure of the *P. aeruginosa* type strains to GA was once again compared to that of AMPs CST and LL-37. No significant differences were seen in the expression of any of the TCS genes, which had been normalised to expression of the housekeeping gene *rpsL* (ΔCt), with any intervention (**Supplementary Data**). For the ΔΔCt, changes compared to untreated cultures, exposure to LL-37 resulted in expression increases >2-fold of the TCS genes *phoP* (4.06 [95%CI 1.26-4.20]) and *pmrA* (5.08 [95%CI 1.91-9.13]) and the modification gene *arnB* (8.55 [95%CI 4.29-11.54]). Exposure to GA and CST did not result in a ΔΔCt of >2-fold increase for any of the genes tested in the type *P. aeruginosa* strains (each of which is sensitive to CST) (**Figure 6**). These results are in keeping with the above Lipid A modification data where only LL-37 stimulates any response from the type *P. aeruginosa* strains.

**Figure 6.**
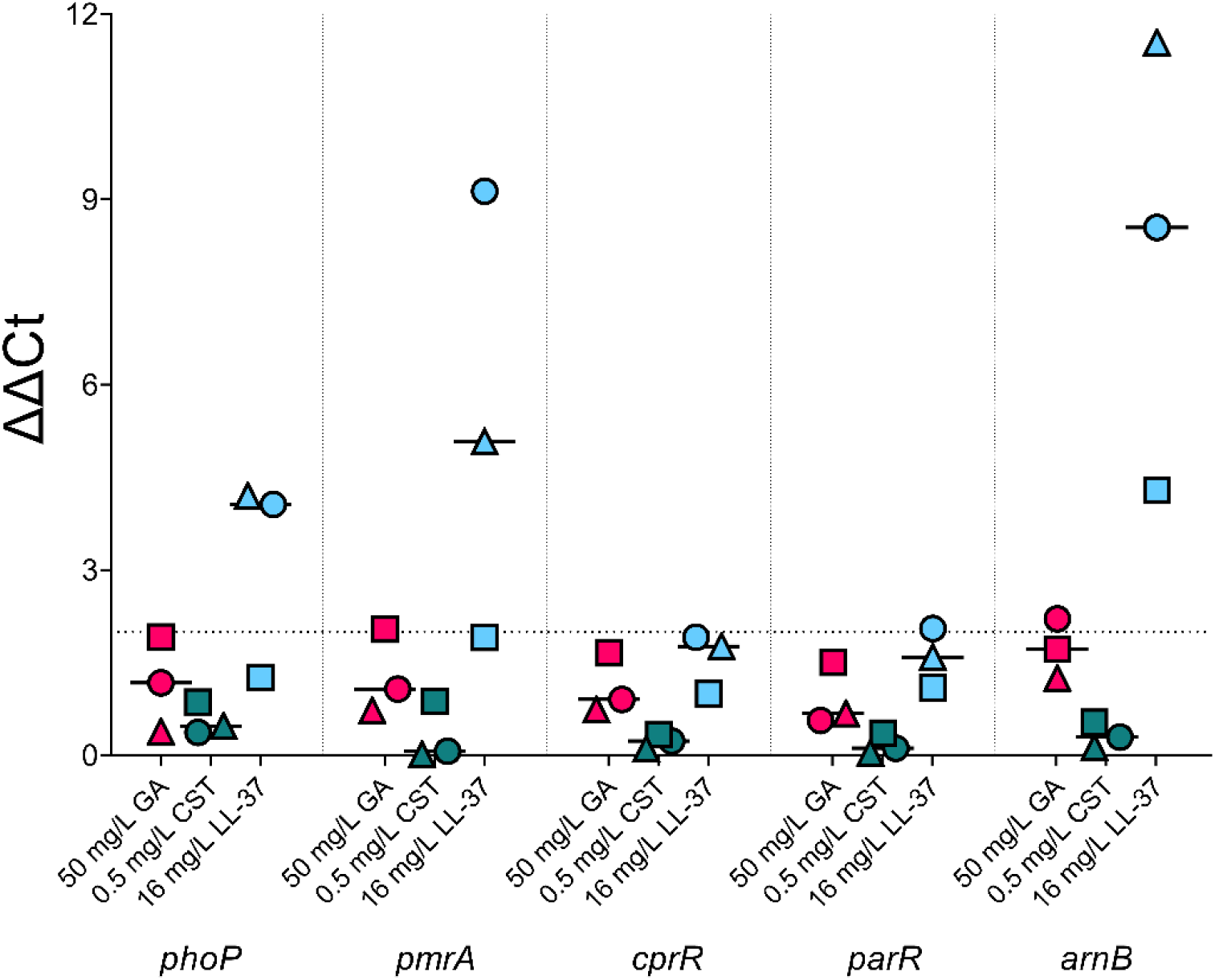
Expression of Two Component System genes phoP, pmrA, cprR and parR and L-Ara4N modification gene arnB of P. aeruginosa strains PAO1, PA14 and PAK after exposure to AMPs GA, CST and LL-37. ΔΔCt results of gene expression, normalised to untreated P. aeruginosa. Only exposure to LL-37 resulted in fold increases in gene expression > 2 with median expression of phoP of 4.06 (95%CI 1.26 - 4.20), pmrA of 5.08 (95%CI 1.91 - 9.13) and arnB of 8.55 (95%CI 4.29 - 11.54). Each point is the median of biological replicates for P. aeruginosa PAO1 (ϒ), PA14 (♦) or PAK (ρ). Line at median.

### Expression of Two Component Systems and *arn* operon genes of clinical *P. aeruginosa* from people with CF after GA exposure

Clinical *P. aeruginosa* isolates from people with CF were next tested for their reaction to GA exposure using the same set of genes as the type strains. Exposure to GA resulted in a significant increase in the expression of genes *pmrA* (p < 0.05) and *arnB* (p < 0.01) compared with untreated cultures (**Supplementary Data**). However, only a modest (< 2-fold) median ΔΔCt change was seen for each (*pmrA* of 1.39 [95%CI 0.82 - 2.52] and *arnB* of 1.55 [95%CI 1.16 - 2.08]) (**Figure 7A**). Change in expression of these two genes after exposure to 50 mg/L GA was also positively correlated (p < 0.005, Spearman r = 0.8 [95%CI 0.36 - 0.95]) (**Figure 7B**).

**Figure 7.**
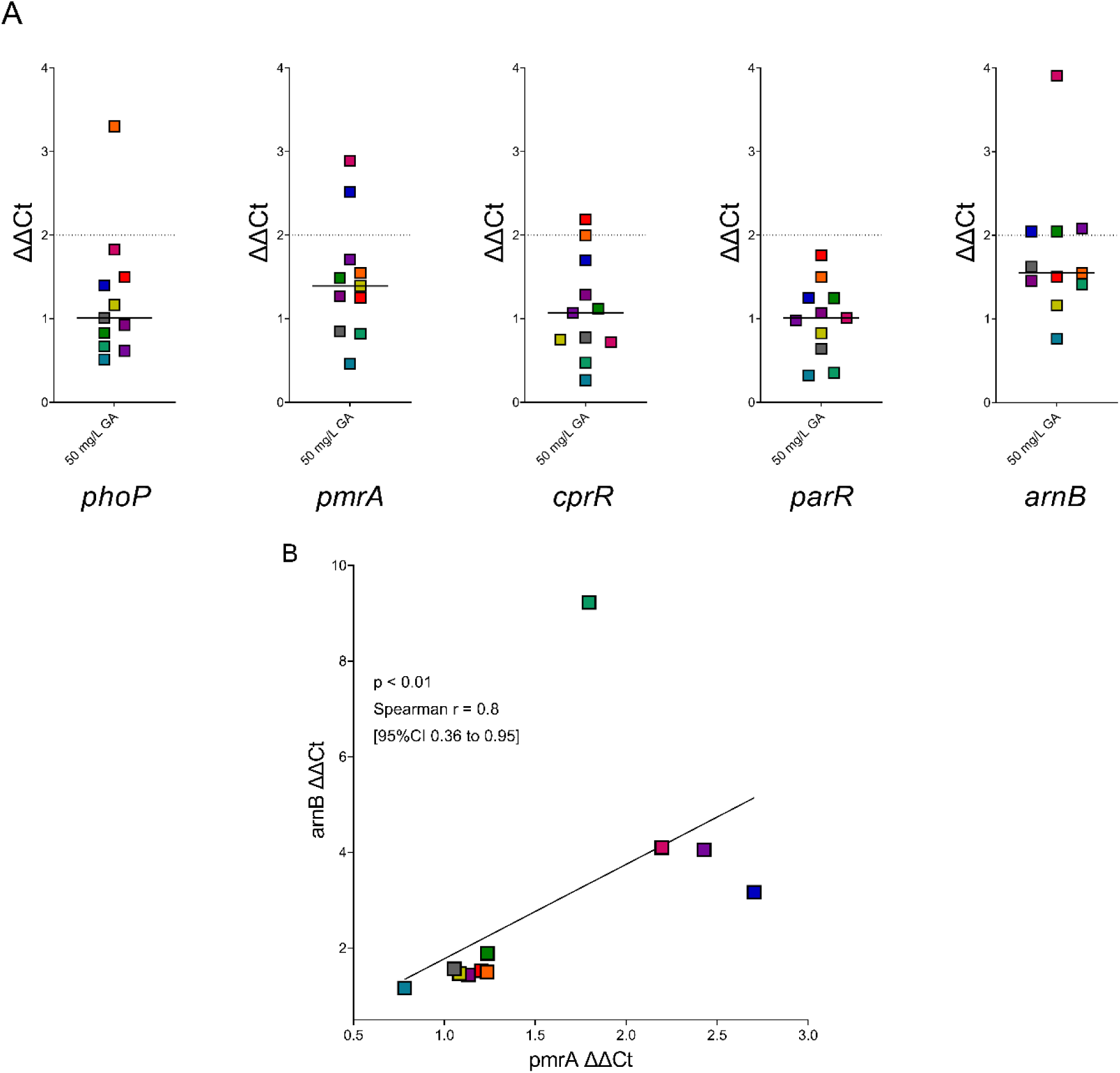
Expression of Two Component System genes phoP, pmrA, cprR and parR and L-Ara4N modification gene arnB of P. aeruginosa clinical strains from people with CF after exposure to GA. **A**. ΔΔCt results of gene expression, normalised to untreated P. aeruginosa for each strain. No gene tested had > 2-fold median increase across the 11 clinical P. aeruginosa tested. On each graph, each point is the median of biological replicates for a clinical P. aeruginosa strain, line at median. **B**. Correlation analysis of ΔΔCt expression levels of pmrA and arnB. Across the clinical strains tested, there was significant positive correlation between the expression of pmrA and arnB after exposure to GA (p = 0.0047, Spearman r = 0.8 [95%CI 0.36 - 0.95]). Strain colour coding and values can be found in Supplementary Material.

The remaining genes tested were not significantly altered by GA in clinical *P. aeruginosa,* when compared to untreated cultures, and each had expression fold changes < 2; *phoP* 1.01 (95%CI 0.62 - 1.83), *cprR* 1.07 (95%CI 0.48 - 2.00) and *parR* 1.01 (95%CI 0.36 - 1.50) (**Figure 7A**). No correlation was seen in the expression levels for any of these TCS genes with expression of *arnB* after exposure of the clinical *P. aeruginosa* strains to GA.

Resistance to CST and other AMPs has been associated with specific, non-synonymous mutations in TCS genes; in true resistance these SNPs frequently lead to inactivation of the sensor genes of TCS(s) which results in constitutive activation of the L-Ara4N modification system encoded by the Arn operon ^84^. To confirm that TCS genes from the 11 clinical strains tested were not identical, the sequences examined for SNPs resulting in amino acid changes. All clinical isolates had at least one amino acid (aa) change to one of the TCS genes examined (compared with PAO1). A total of 24 aa changes were identified in clinical isolates TCS genes ranging from 1 to 11 (**Table 1, Supplementary Table 3**). The strain with the highest number of aa changes (GA_899) had aa changes in the sequence of PmrB which have been associated with an intermediate level of CST resistance in clinical *P. aeruginosa* previously but gene expression was < 2-fold for all genes tested in this strain ^85^. (Expression of the ColSR system was not measured in this study and no amino acid changes were found in any strain in this system).

**Table 1.**
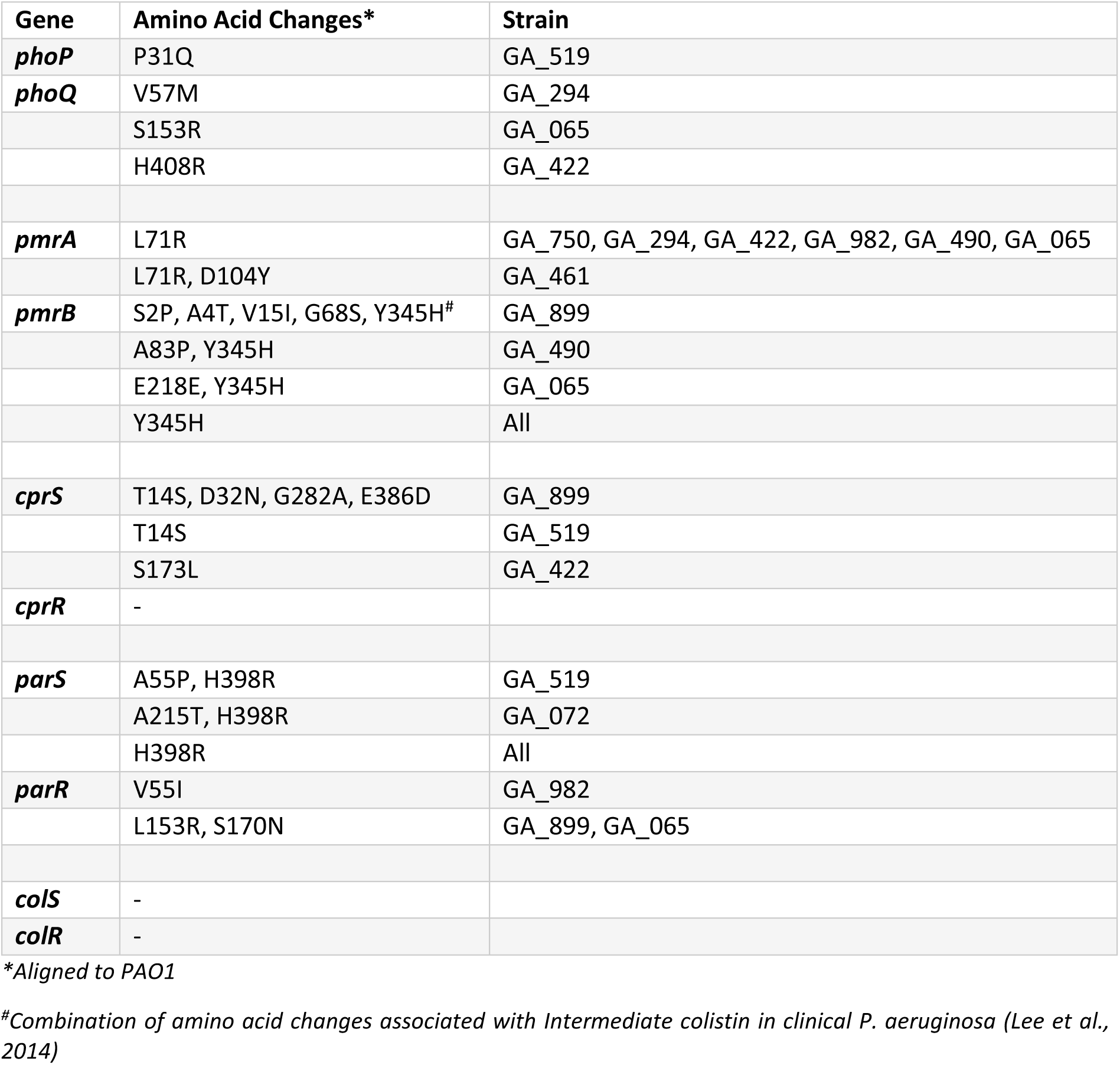
Non-synonymous changes found in the gene sequences of sensory components of Two Component Systems of 11 clinical P. aeruginosa strains from cystic fibrosis. For gene accession numbers see Supplementary Table 3.

## DISCUSSION

The evidence from our previous work was that GA acts on *P. aeruginosa* inner and outer cell membranes, damaging both and permeabilising the cell, and it was therefore of interest to test whether electrostatic interactions with LPS were the point of contact between GA and the bacterial cell, as seen with many other cationic AMPs ^50,51,61,86^. Here we have confirmed the binding of GA and *P. aeruginosa* LPS to each other via both LPS quantification and GA activity. GA sequesters and neutralises *P. aeruginosa* LPS and, conversely, LPS can sequester GA. These results indicate that the LPS on the surface of *P. aeruginosa* is a cellular target for GA, via its cationic properties allowing binding to the bacterial cell and resulting in its membrane perturbing effects.

With this information it was therefore necessary to investigate the significance for GA activity of GA-LPS interactions, particularly given the salience of free-LPS in CF sputum ^39,41,56,87^. The results here for membrane perturbation assays further demonstrate interaction between GA and *P. aeruginosa* LPS; the presence of a supraphysiological concentration of LPS significantly reduced GA disruption of the OM and depolarisation of the CM. However, a CF physiological LPS concentration (0.02 mg/mL) has less impact: across the three membrane assays and the three *P. aeruginosa* strains tested, only OM disruption was reduced. Divalent cations of Mg^2+^ were also protective against previously reported OM disruption and CE permeabilisation activities of GA ^51^. This is further confirmation of our earlier observation of GA and LPS binding and of competition between GA and Mg^2+^ for LPS binding, which indicates that the interaction is driven by their opposing charges, a feature of AMP activity ^31^. While we recognise testing these elements in isolation of each other may not reflect the full complexity of the CF lung environment – which may be more detrimental to GA activity than each element alone – the elements tested here are also known to interact with and/or sequester each other and may exert a less confounding effect on GA due to their competitive binding with each other ^31,46^.

We note that the protective effect of Mg^2+^ cations against GA activity was seen for the OM and CE, but absent for the CM. This may indicate that the activity of GA against *P. aeruginosa* cells is multimodal rather than via one distinct mechanism. GA is a random peptide with huge heterogeneity (formed of up to 10^30^ possible peptides), has the ability to adopt more than one conformation in solution and to oligomerise, therefore evidence of activity against the bacterial cell via more than one mechanism is perhaps not surprising ^54,55,88^.

We were also interested in investigating any interactions between GA and eDNA. As with LPS, eDNA is an important component of CF sputum; it is negatively charged, important in *P. aeruginosa* infection and has been shown to be a factor in AMP resistance in bacteria ^46,47,78,89^. There were also previous indications of DNA aggregation by GA ^50^. In common with our observations for LPS, here we have shown that there is direct interaction between GA and eDNA; quantifiable eDNA (1 and 10 mg/mL) was reduced by GA in a dose-dependent manner, significantly so at GA ≥ 25 mg/L. At a physiologically relevant concentration of eDNA, the activity of GA was not impaired ^58^.

With confirmation of GA interactions with LPS as a mechanism of GA activity, this leads to the question of whether or not *P. aeruginosa* strains would mount a defence against the targeting of its LPS by GA. Resistance to AMPs, such as CST, is associated with the modification of the Lipid A portion of LPS; the addition of positively charged moieties of various kinds to neutralise the charge of the bacterial cell and limit AMP binding ^63,90,91^. The modification of the Lipid A component of LPS was tested using MALDI-TOF analysis on which we saw no significant changes in any of the Lipid A additions tested across the type *P. aeruginosa* strains. These results show that GA exposure does not result in a significant response being mounted by the type strains in response to GA activity, in the manner commonly described for other AMPs. For further details on the reaction to AMP exposures, the expression of genes involved in TCSs known to detect AMPs in *P. aeruginosa* (*phoPQ*, *pmrAB*, *cprSR* and *parSR*) and a gene of the *arn* operon were tested. These TCSs feed information to the Arn operon which controls the modification of LPS with L-Ara4N, as protection against AMP attack. We found no significant changes in gene expression levels across the type strains of *P. aeruginosa* due to AMP exposure, including GA. Only the human cathelicidin LL-37 resulted in median ΔΔCt > 2- fold across the type strains which was the case for *phoP*, *pmrA* and *arnB*.

In clinical *P. aeruginosa* strains, no significant increases in Lipid A additions were seen after GA exposure. Expression of genes *pmrA* and *arnB* were increased in clinical strains due to GA exposure, but in neither case > 2-fold greater than untreated bacteria. There was a correlation between the expression of the two significantly increased genes indicating that, even at the low level of the effect recorded, sensing of and reaction to GA by clinical *P. aeruginosa* is taking place via the well documented cascade of PmrAB to the Arn operon ^36,85^. This suggests that the clinical *P. aeruginosa* did not fail to detect the presence of GA even if this did not lead to a notable fold increase in gene expression nor Lipid A modification by addition of L-Ara4N in clinical strains. This is despite L-Ara4N addition being the process which the Arn operon mediates. Neither was an increase in any of the other Lipid A modification types tested here seen due to GA exposure.

The search for solutions to the global AMR crisis has resulted in a renewed and increased interest in the utility of AMPs ^20^. While issues of cytotoxicity at effective, antibacterial doses has limited the number of AMPs transferring to the clinic so far – unlikely to be an issue for the already clinically used GA – many AMPs are under investigation as new antimicrobials and AMPs have several advantages over conventional antibiotics and other novel therapies ^7,8,20^. Chief among these is the low rate of resistance generation resulting from AMP exposure; AMPs have been shown to be less likely to produce resistance, to result in a lower recombination rate than antibiotics and CST resistance took longer to emerge than resistance to other antibiotics ^24,33,92^. While concerns have been raised about cross-resistance between AMPs – due to their similar mechanisms of action – we did not note any crossover between the reaction of clinical *P. aeruginosa* to GA and their CST-sensitivity phenotypes, in our short term exposures ^93^. The clinical strain with the strongest reaction to GA in its LPS modification (GA_899) was designated CST-sensitive clinically and the CST-resistant strain in the panel (GA_422) did not mount a strong LPS modification reaction to GA exposure. Encouragingly, overall the results presented here for GA indicate that *P. aeruginosa* does not mount a strong response to GA activity in a manner seen for exposure to other AMPs.

Moreover, our previous work proposes a role for GA less as a direct acting antibiotic and more as an antibiotic adjuvant to be given in combination with the aminoglycoside tobramycin, with which we demonstrated to have substantial synergy ^51^. The non-bactericidal nature of GA at a synergistic concentration and the absence of a resistance response in the results presented here both suggest reduced selective pressures for AMR development by *P. aeruginosa* strains in response to GA exposure. As mentioned, GA is a random peptide and random peptides are currently being researched as “resistance proof” antimicrobials with synergy between AMPs being well documented and resistance evolution having been shown to be less common for AMP combinations/cocktails than single AMPs or conventional antibiotics ^94–98^. A co-treatment strategy with a conventional antibiotic further alleviates concerns for the future generation of resistance with combinations and synergy less likely to result in antimicrobial resistance and loss of utility (while also rescuing the efficacy of tobramycin in the case of GA)^96,99–102^.

The results presented here point to further potential ancillary benefits of GA – on top of the known advantages of antibiotic synergy, little bacterial response and its immunomodulatory properties – with evidence that both *P. aeruginosa* LPS and eDNA were stably bound and sequestered by GA. As well as being highly immunogenic in infection generally, free-LPS is particularly important in CF lung infections ^40,103^. In common with other areas of the body, LPS is pro-inflammatory in CF sputum and contributes to NETosis, a key driver of cycles of inflammation and bacterial infection in CF ^43–45^. Free-LPS has also been shown to exacerbate the CFTR defect and to contribute to longer-term lung damage ^104,105^. NETosis can also lead to an increase in the eDNA content of CF sputum which is detrimental to the lung in a number of ways; eDNA is a vital component of *P. aeruginosa* biofilms, increases resistance to AMPs and contributes to sputum acidification resulting in increased AMR and viscosity ^47,78,89^. Therefore, neutralisation of either/both of these sputum components could be of benefit in reducing inflammation and cycles within the CF lung which can promote *P. aeruginosa* persistence ^48,49,76,77,86^. Both CST and LL-37 have been investigated as LPS neutralisers, to beneficial effect, as well as other AMPs ^48,75,76,106^.

Gram-negative LPS is a crucial point of contact in host-pathogen interactions. LPS is detected by host immune systems as a marker of infection and LPS is targeted as a binding site for host defence AMPs but LPS and its modification also provide bacteria with a defence against antimicrobials. In CF, this is further complicated by the presence of free-LPS in the airway. Here, we have demonstrated that LPS plays a role in the activity of GA but the latter was not extensively inhibited by free-LPS when present in the background. We also showed that, in both type strains and isolates from people with CF, GA did not trigger a strong response in LPS modification by *P. aeruginosa*, either at the genetic or physical level, to defend its LPS against GA. This appears compatible with the fact that GA is not exerting strong selective pressure on systems frequently associated with AMP resistance. Interest in research into random peptide cocktails is increasing with the aim of producing “resistance proof” antimicrobials ^94–96^. These results add to the previously published evidence for GA as an AMP with strong suitability for repurposing and with a strong, longstanding safety profile could be a forerunner of random peptides as treatments for infection.

### Data Availability

The data that support the findings of this study are available from the corresponding author upon reasonable request. Gene accession numbers can be found in Supplementary Table 3.

## ACKNOWLEDGEMENTS

Authors would like to thank the Quadram Institute Biosciences Core Sequencing and Core Bioinformatics teams their assistance in this project.

