## Supplementary Tables for "Antimicrobial peptide glatiramer acetate targets *Pseudomonas aeruginosa* lipopolysaccharides to breach membranes without altering lipopolysaccharide modification"

Supplementary Table 1. Clinical P. aeruginosa isolates used in this study.

| Strain |  | Sample Type | Mucoidy | Multidrug Resistance | CST | TOB | GA/TOB Synergy* |
| --- | --- | --- | --- | --- | --- | --- | --- |
| GA_750 |  | Spontaneous Sputum | Mucoid |  | S | S | ✓ |
| GA_899 |  | Spontaneous Sputum |  |  | S | S |  |
| GA_550 |  | Spontaneous Sputum |  |  | S | S |  |
| GA_294 |  | Spontaneous Sputum | Mucoid |  | S | S |  |
| GA_982 |  | Spontaneous Sputum |  |  | S | S | ✓ |
| GA_422 |  | Spontaneous Sputum |  |  | R | S | ✓ |
| GA_519 |  | Spontaneous Sputum | Mucoid |  | S | R | ✓ |
| GA_461 |  | Cough Swab |  | MDR | S | R |  |
| GA_490 |  | Spontaneous Sputum |  | MDR | S | R | ✓ |
| GA_065 |  | Spontaneous Sputum |  |  | S | R |  |
| GA_072 |  | Spontaneous Sputum |  |  | S | R | ✓ |

*S – sensitive; R – resistant*

*MDR – multidrug resistant i.e. resistant to three or more antibiotics from different classes ^1,2^*

**characterised in Murphy et al., 2022 ^3^*

Supplementary Table 2. Mass to charge (m/z) ratios of unmodified Native Lipid A and each type of modified Lipid A measured by MALDI-TOF.

|  | m/z |
| --- | --- |
| Native Lipid A | 1404 1447 1462 |
| Phosphate | 1484 1527 1542 1654 1697 1712 1722 1765 1780 |
| C10:30OH | 1494 1537 1552 1574 1617 1632 1654 1697 1705 1712 1748 1763 1812 1836 1855 1870 1879 1894 |
| Palmitate | 1562 1605 1620 1642 1685 1700 1722 1765 1773 1780 1812 1816 1831 1855 1870 1904 1947 1962 |
| L-Ara4N | 1535 1578 1593 1666 1705 1709 1724 1748 1763 1773 1816 1831 1836 1879 1894 1904 1947 1962 |

Supplementary Table 3. GenBank Accession Numbers.

|  | *phoP* | *phoQ* | *pmrA* | *pmrB* | *cprS* | *cprR* | *parS* | *parR* |
| --- | --- | --- | --- | --- | --- | --- | --- | --- |
| GA_750 | OR023699 | OR023710 | OR023677 | OR023688 | OR023633 | OR023644 | OR023655 | OR023666 |
| GA_899 | OR023700 | OR023711 | OR023678 | OR023689 | OR023634 | OR023645 | OR023656 | OR023667 |
| GA_550 | OR023698 | OR023709 | OR023676 | OR023687 | OR023632 | OR023643 | OR023654 | OR023665 |
| GA_294 | OR023693 | OR023704 | OR023671 | OR023682 | OR023627 | OR023638 | OR023649 | OR023660 |
| GA_982 | OR023701 | OR023712 | OR023679 | OR023690 | OR023635 | OR023646 | OR023657 | OR023668 |
| GA_422 | OR023694 | OR023705 | OR023672 | OR023683 | OR023628 | OR023639 | OR023650 | OR023661 |
| GA_519 | OR023697 | OR023708 | OR023675 | OR023686 | OR023631 | OR023642 | OR023653 | OR023664 |
| GA_461 | OR023695 | OR023706 | OR023673 | OR023684 | OR023629 | OR023640 | OR023651 | OR023662 |
| GA_490 | OR023696 | OR023707 | OR023674 | OR023685 | OR023630 | OR023641 | OR023652 | OR023663 |
| GA_065 | OR023691 | OR023702 | OR023669 | OR023680 | OR023625 | OR023636 | OR023647 | OR023658 |
| GA_072 | OR023692 | OR023703 | OR023670 | OR023681 | OR023626 | OR023637 | OR023648 | OR023659 |

1.	Santajit, S. & Indrawattana, N. Mechanisms of Antimicrobial Resistance in ESKAPE Pathogens. *Biomed Res. Int.* **2016**, (2016).

2.	Magiorakos, A. P. *et al.* Multidrug-resistant, extensively drug-resistant and pandrug-resistant bacteria: An international expert proposal for interim standard definitions for acquired resistance. *Clin. Microbiol. Infect.* **18**, 268–281 (2012).

3.	Murphy, R. A. *et al.* Synergistic Activity of Repurposed Peptide Drug Glatiramer Acetate with Tobramycin against Cystic Fibrosis Pseudomonas aeruginosa. *Microbiol. Spectr.* e0081322 (2022) doi:10.1128/spectrum.00813-22.
